# makeTCR: A Modular Platform for Rapid, Flexible, Scalable, Single-Step T Cell Receptor Synthesis

**DOI:** 10.1101/2025.04.27.647198

**Authors:** M. Hamberger, M-T. Neuhoff, S.V. Pietrantonio, T. Boschert, C. Maldonado Torres, A. Errerd, C. L. Tan, J. M. Lindner, M. Platten, E. W. Green

**Author notes:** Contributed equally, Correspondence. **Author Contributions (<u>CRediT Taxonomy</u>)** - MH: Methodology, Investigation, Writing – review & editing - M-TN: Investigation, Methodology, Formal analysis, Writing – original draft, Visualization - SVP: Methodology, Software - TB: Methodology, Investigation, Formal analysis, Writing – review & editing - CMT: Investigation - AE: Investigation, Visualization - CLT: Methodology - JML: Resources, Writing – review & editing - MP: Funding acquisition, Resources, Supervision, Writing – review & editing - EWP: Conceptualization, Investigation, Software, Writing – original draft, Writing – review & editing, Project Administration, Supervision, Funding acquisition. **Competing interests**: MP, CLT and EWG hold patents and PCT applications describing methods to identify tumor-reactive T cells, and are founders or employees of Tcelltech GmbH. JML is an employee of BioMed X GmbH.

## Abstract

Personalised cell therapies utilising T cell receptors (TCRs) show tremendous clinical promise, though TCR synthesis and validation techniques lag behind our ability to sequence TCR repertoires. To address this gap, we developed makeTCR: a flexible modular, scalable TCR cloning system that enables single-step, 100% fidelity assembly of αβ and γδ TCRs into commonly used expression vectors. By implementing cell-free manufacturing, makeTCR enables patient-derived TCRs to be functionally validated within 48 hours. We provide an open-source, easily extensible, web-based graphical platform that integrates existing tools to simplify and standardise the manufacture of TCRs across all scales of synthesis.

## Introduction

Human adaptive immunity depends on T cells, which respond to immune challenge by recognising short peptides presented on major histocompatibility complex proteins (pMHC) by means of their unique heterodimeric T cell receptor (TCR). In humans, the process of V(D)J recombination ensures enormous TCR sequence diversity; at any given time a person has around 10^10^ unique TCR clonotypes, allowing for the recognition of a vast range of antigens – including auto antigens in the case of autoimmune disease. TCRs recognising recurrent tumor antigens such as MAGE-A4 ^1^ or PRAME ^2^ have been used to genetically engineer autologous T cells to create cellular immunotherapies such as the FDA approved afamitresgene autoleucel. Unfortunately, the costs of identifying TCRs recognising a specific pMHC complex remain high, with the result that most TCR immunotherapies in development target recurrent tumor epitopes presented by the HLA-A*02:01 allele found in about half of Caucasians.

For cancer patients lacking HLA-A*02:01 expression, or whose tumors do not express an antigen targeted by an ‘off-the-shelf’ TCR therapy, TCRs reactive to patient-specific epitopes can be identified - in some cases resulting in remarkable remissions ^3^. Such clinical outcomes raise the hope that personalised autologous TCR T cell therapies may offer clinical benefit for a broad range of patients, however the median vein-to-vein time for creating such a personalised TCR therapy exceeds six months ^4^. Recently we and others have shown that the first step of this process – the identification of tumor-reactive TCRs – can be accelerated by identifying these T cells within the TIL population on the basis of single cell sequencing data using gene expression signatures ^5–9^ or machine learning classifiers ^10,11^. However, the manufacture of candidate TCRs remains complex and costly, even when using pre-cloned libraries of TCR V genes with synthetic CDR3-J sequences using either Golden Gate ^12^ or Gibson assembly ^13,14^. Previous TCR cloning approaches were limited by requiring multi-step processes that supported only the manufacture of human αβ TCRs within a single expression vector utilising a single TCR constant region, and also lacked software tools to automate the process of cloning TCRs from TCR repertoire data.

Here we present makeTCR, a modular, Golden Gate-based TCR cloning system that enables single-step, 100% fidelity cloning of both human and mouse, αβ and γδ TCRs into an extensive collection of modular TCR expression vectors and engineered TCR constant domains.

makeTCR’s flexibility allows researchers to generate vectors compatible with their existing workflows, whilst benefitting from reduced costs, complexity and turnaround time and increased flexibility. We show that the makeTCR platform can generate entirely novel, patient-derived TCRs in 24 hours, allowing candidate therapeutic TCRs to be validated and prioritised prior to therapy.

To maximise the utility of makeTCR we developed a freely available, open source, graphical web-based companion software platform to automate and simplify the generation of TCRs and HLA-appropriate controls from TCR repertoires and databases that scales from cloning individual TCRs to whole repertoires. Lastly, we show that users can easily propagate hundreds of precloned modules using rolling circle amplification (RCA) in plate-based formats without deleterious effects on TCR assembly fidelity or error rate, improving the useability, cost, and ultimately throughput of TCR synthesis.

## Results

### Flexible, high-efficiency modular TCR assembly

We designed a series of mutually orthogonal Golden Gate overlaps to enable BsmBI mediated assembly of αβ and γδ TCRs by combining four pre-cloned modules encoding the germline variable and constant domains of a TCR with a vector backbone and *de novo* synthesised CDR3-TR*J fragments (Figure 1a). By utilising silent codon mutagenesis we generated a set of overlaps between modules predicted to result in 100% fidelity assembly for αβ and γδ TCRs (Figure 1b-d, Supplemental Figure 1). We used these overlaps to split the blue chromoprotein amilCP ^15^ into six modules, and experimentally confirmed assembly fidelity to be 99.9% using blue/white colony screening in *E.coli* (Supplemental Figure 2a). We used this blue/white assay to rapidly test variant Golden Gate thermocycling protocols, developing an optimised protocol that doubled assembly yield relative to the manufacturer’s recommendations (Figure 1e), allowing us to miniaturise TCR assembly reactions without compromising yield.

**Figure 1:**
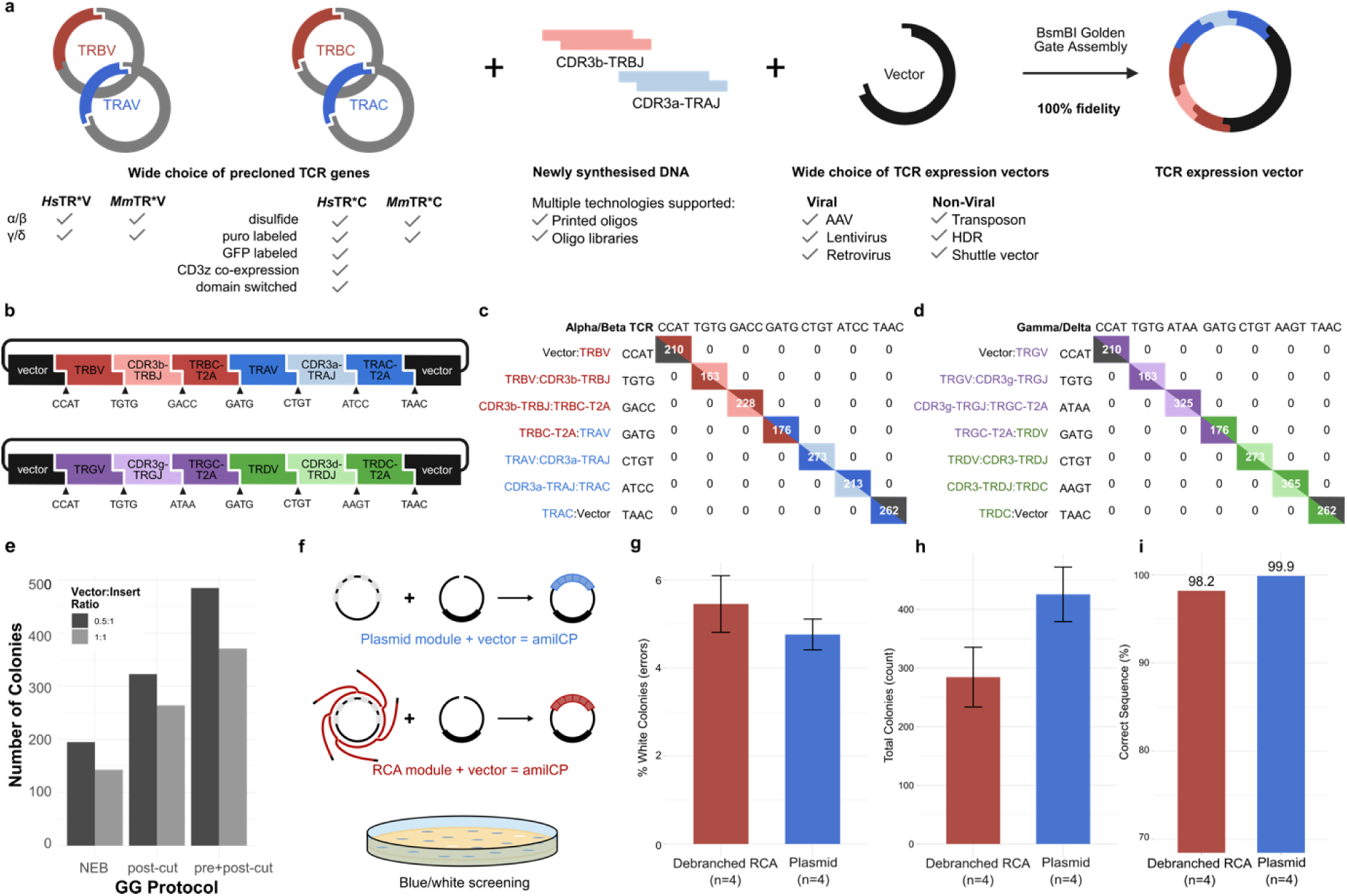
Modular, high fidelity Golden Gate TCR assembly system using makeTCR. **a**, Schematic for the makeTCR system of BsmBI mediated modular Golden Gate assembly of TCRs using pre-cloned germline TR*V and TR*C genes, newly synthesised DNA oligonucleotides encoding the CDR3-TR*J regions and a panel of compatible TCR expression vectors. **b,** schematic of an assembled polycistronic αβ or γδ TCR expression construct. **c**;**d**, ligation fidelity tables showing 100% expected fidelity when assembling αβ or γδ TCRs from makeTCR encoded modules. **e**, modular assembly of the amilCP expression cassettes was used to test variant BsmBI mediated Golden Gate assembly thermocycling protocols; a variant protocol with pre- and post-assembly thermal soak steps resulted in a twofold increase in yield (number of blue colonies). **f**, Experimental setup comparing the efficiency and fidelity of Golden Gate assembly using either plasmid or RCA amplified modules encoding the amilCP chromoprotein **g**, No significant difference in the overall fidelity (number of blue vs white colonies) was found when using plasmid or RCA amplified input (P=0.391, Welch Two Sample t-test, n=4 replicates per template type). **h**, Golden Gate assembly using RCA encoded modules resulted in a non-significant drop in yield vs using plasmid encoded modules (P=0.087, Welch Two Sample t-test, n=4 replicates per condition. **i**, Nanopore sequencing of completed Golden Gate assembly reaction product shows that RCA amplified plasmids exhibit a minor increase in number of SNPs (99.97% perfect reads vs 98.18% respectively).

We then generated plasmids encoding the human and murine germline alpha, beta, gamma and delta variable and constant gene sequences flanked by BsmBI recognition sites (Supplemental Table 1). To maximise utility as a ‘drop-in’ replacement for existing TCR cloning methods we also synthesised a range of commonly used engineered TCR constant regions including variants that limit cross-pairing with endogenously expressed TCRs by incorporating disulfide mutations ^16^ or switched transmembrane domains ^17^, variants that increase the amount of surface-expressed TCR by co-expressing CD3z ^18^, and variants including selection markers mStaygold2 or puromycin N-acetyltransferase to facilitate enrichment or selection of TCR transgenic cells.

### A diverse collection of Golden Gate compatible expression vectors

TCRs have been delivered to cells using a panoply of expression vectors including lentivirus, retrovirus, adeno-associated virus (AAV), transposons, and CRISPR/Cas9 mediated knock-in to the genome – typically at the *TRAC* or the *AAVS1* safe harbour loci. We therefore engineered makeTCR compatible versions of each expression vector backbone (Supplemental Figure 3), including homology directed repair (HDR) cassettes that allow for TCR knock-in to the Human *TRAC* locus using either exon targeting ^19^ or intron targeting guides ^20^. In the case of the commonly used pMXS retroviral vector we used structure-guided engineering to deplete BsmBI sites while preserving secondary structure loops in the MMLV Ψ packaging domain. We also created a makeTCR compatible vector containing attL sites, allowing TCRs to be shuttled into existing Gateway^TM^ destination vectors using LR cloning.

To ensure high-fidelity cloning of TCRs we engineered all expression vector backbones to incorporate unique SwaI sites to enable vector linearisation (reducing input carryover during vector assembly reactions), and made the vectors cross-compatible with the T-CRAFT high-throughput cloning system by adding appropriate SapI sites ^21^. This resulted in a diverse library of Golden Gate compatible vectors supporting high fidelity cloning of TCRs (or other cargoes) to match all commonly used expression vectors, allowing researchers to use makeTCR to complement their existing TCR expression systems.

### Simple propagation of cloning modules using rolling circle amplification

One limitation of modular cloning systems is the requirement to maintain large numbers of modules; plasmid DNA preparation from overnight bacterial cultures is a labour- and space-intensive process that is challenging to automate. In contrast, randomly primed, Phi29-mediated rolling circle amplification (RCA) is a simple, cost-effective, high-fidelity method to propagate circular DNA at concentrations similar to a bacterial midiprep using an isothermal reaction that is compatible with plate-based automation ^22^. Using our modular amilCP cloning system we could show that assembling amilCP from RCA-amplified input modules resulted in equivalent assembly accuracy (fraction white colonies, P=0.391, Figure 1f,g) and a comparable yield (total number of colonies, P=0.087, Figure 1h) compared to using plasmid encoded modules. We used nanopore sequencing to assess the fidelity of assembled products and detected a small increase in SNPs when using RCA amplified templates (99.18% perfectly aligning reads vs 99.97% using plasmid encoded templates, Figure 1i), confirming previous observations ^23^. Given the ease of maintaining module collections by RCA, and the minor increase in sequence error well below the limits of detection for functional TCR validation assays, we performed all further TCR assembly experiments using RCA amplified TR*V and TR*C modules.

### Patient-derived TCR prototyping within 48 hours using makeTCR cloning

We previously showed that personalised T cell immunotherapies can be accelerated by using a machine-learning classifier that identifies tumor-reactive TCRs in an antigen-agnostic fashion amongst tumor infiltrating lymphocytes (TILs) using single cell sequencing data ^10^. We reasoned that combining rapid TCR identification (predicTCR) and manufacture (makeTCR) to prototype TCRs prior to use in personalised transgenic T cell immunotherapies would likely generate a better TCR product. By using pre-cloned modules makeTCR requires only *de novo* DNA synthesis of the relatively short (<90bp) CDR3a-TRAJ and CDR3b-TRBJ regions; regions that can be generated rapidly in-house using benchtop oligonucleotide synthesisers. We therefore predicted additional tumor-reactive TCRs from our previously described ‘BT21’ melanoma brain metastasis (Supplemental Table 2) and optimised a cell-free TCR manufacturing strategy that allowed us to generate messenger RNA (mRNA) encoding these TCRs within 24 hours (Figure 2a) – a process that would take over a week using existing approaches. This rapid manufacturing strategy did not reduce overall TCR quality: Nanopore sequencing showed that over 90% of reads for each TCR were free of sequence errors (Figure 2b), and transfecting candidate TCR mRNA into TCR-deficient Jurkat cells resulted in high levels of the transgenic TCR in the cell membrane (52.0%-98.1% mTCRb^+^ expression, Figure 2c). However, transfection of the same mRNA into healthy donor T cells resulted in a variable, highly repeatable presentation of each TCR ranging from 0.1 % to 32.5% mTCRb^+^ cells (Figure 2c). As candidate TCRs were cloned as chimeric TCRs utilising murine constant domains that do not pair with endogenous human constant domains, this likely reflects the differential affinity of each TCR for limited CD3 molecules in the endoplasmic reticulum during maturation ^18^, with some transgenic TCRs outcompeted by the endogenous TCRs. For three dominantly expressed transgenic TCRs we could confirm reactivity against the BT21 tumor cell line (Figure 2d). These results underscore the importance of functionally prototyping TCRs when identifying the best candidates for personalised cell therapy, especially when not ablating the endogenous TCR, and show that such assays can be completed within 48 hours.

**Figure 2:**
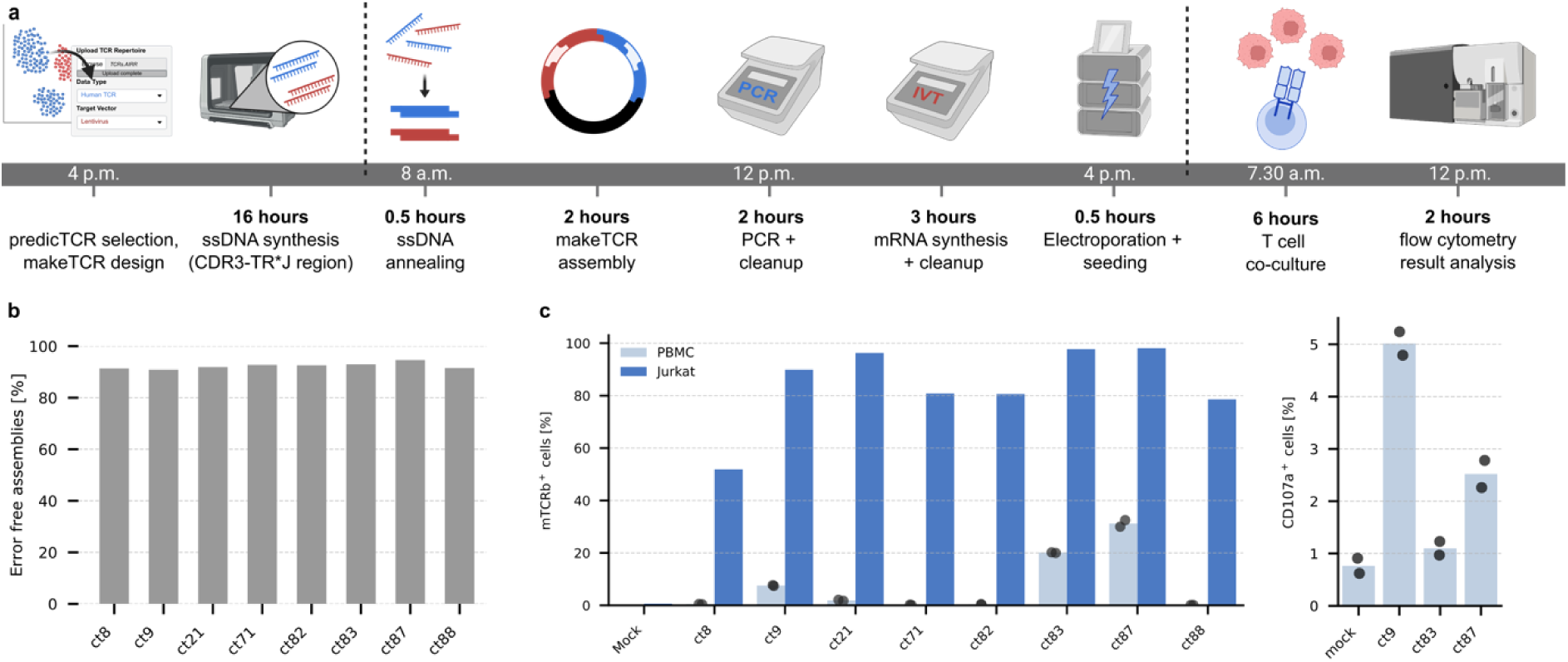
Rapid makeTCR mediated TCR Assembly and TCR Prototyping. **a**, Schematic diagram of rapid, cell-free TCR manufacture and testing using an on-site enzymatic oligonucleotide synthesizer and makeTCR modules to generate a polycistronic mRNA for electroporation into T cells. **b**, Nanopore sequencing of eight TCR clonotypes from BT21 TILs generated using the rapid makeTCR synthesis approach showed >90% error free reads. **c**, Electroporation of IVT generated chimeric TCR mRNA resulted in generally high expression in TCR deficient Jurkat cells (% cells mTCRb^+^), with previously tested clonotypes showing similar patterns as in Tan et. al. (weak expression of ct8, strong expression of ct9). Short term expression of the same mRNA in primary PBMCs lead to heterogeneous expression (ranging from 0.1 to 32.5% mTCRb+), as some transgenic TCRs are outcompeted for access to the CD3 pool. **d**, Co-culture of transgenic REP T cells with the BT21 tumor cell line confirmed the predicted anti-tumor reactivity of strongly expressed TCR clonotypes ct84 and ct87, as well as the ct9 positive control, as assessed by expression of the early activation marker CD107a.

### makeTCR mediated manufacture of large synthetic TCR libraries

Given the increasing demand for large numbers of synthetic TCRs for both basic and translational research, we developed a variant of makeTCR to support low-cost TCR synthesis using oligonucleotide pools (Figure 3a). We encoded each TCR on a 250 bp oligonucleotide containing the TCR’s unique combination of CDR3a-TRAJ and CDR3b-TRBJ sequences bracketed by BsmBI sites, a length-normalizing filler sequence, and a unique pair of mutually orthogonal primers ^24^(Supplemental Figure 4a,b). This design allows individual TCRs to be amplified from an oligonucleotide pool by PCR, a process that can easily be adapted to 96 or 384 well format utilising liquid handling systems. We reasoned that using 96 demultiplexing primers to simplify integration with liquid handling systems allowed for 4,560 unique primer pairs (Figure 3b); at this scale each oligonucleotide can be synthesised for less than €1, resulting in an overall per-TCR manufacturing cost of less than €3.50 (Supplemental Figure 4c) – well below the cost of conventional TCR cloning (€80-200).

**Figure 3:**
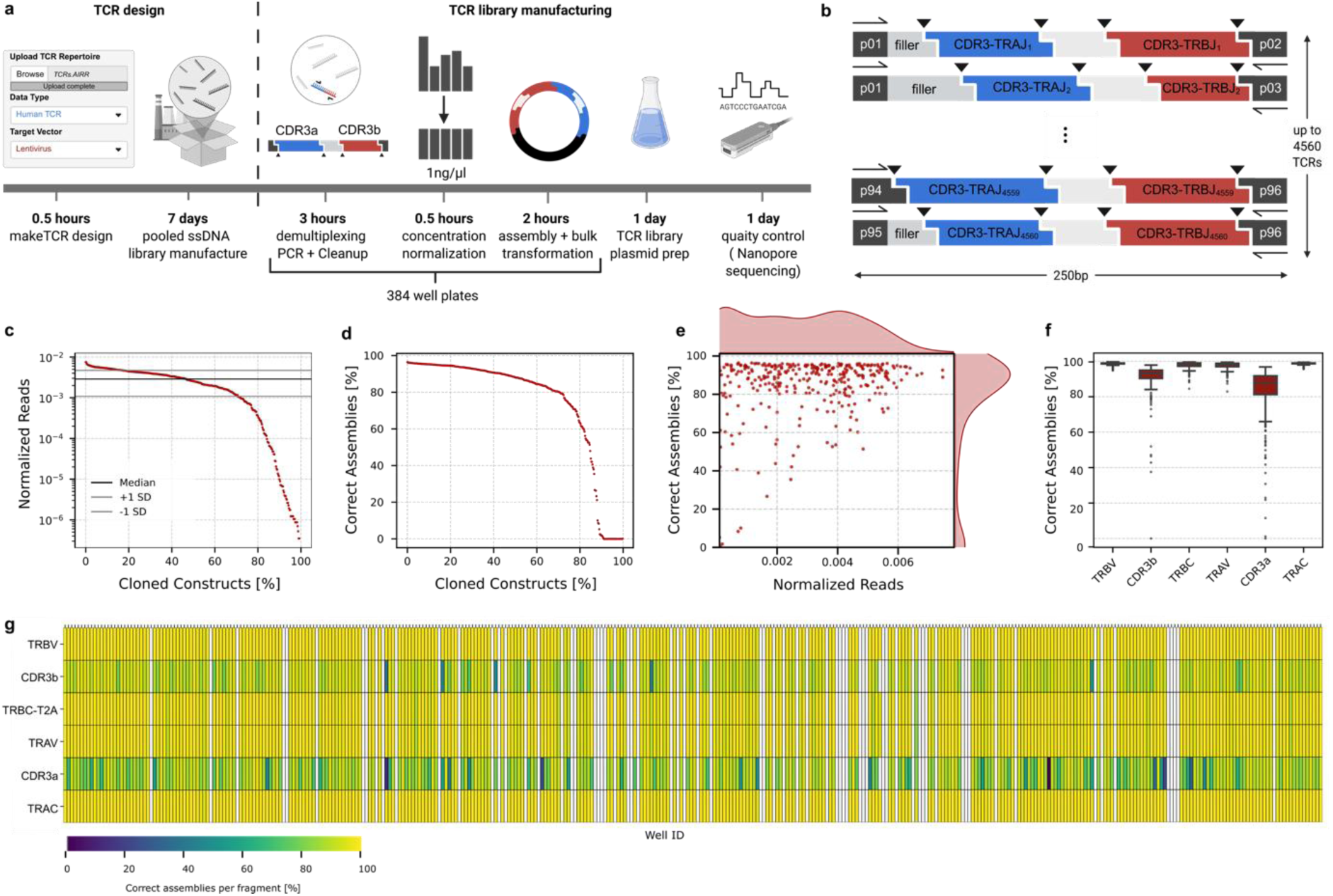
High-Throughput, Oligo-pool mediated makeTCR Assembly. **a**, Schematic illustrating the use of makeTCR to manufacture TCRs in 384 well plate format by demultiplexing pooled ssDNA oligonucleotides. CDR3a-TRAJ and CDR3b-TRBJ pairs are encoded as 250nt oligonucleotides flanked by a unique pair of mutually orthogonal primer binding sites. Oligos are synthesised in a pooled format, demultiplexed, cleaned and normalised in 384 well plates, before being assembled in miniaturised Golden Gate reactions with appropriate TR*V, TR*C and vector backbone modules. Assembled TCRs are pooled and transformed into *E. coli* to generate a TCR library ready for NGS quality control ready for screening as previously described ^36^. **b**, schematic of oligos format using filler sequences to normalise oligo length to 250 nt. 96 unique primers allow up to 4,560 oligos to be individually amplified from the oligo pool. **c**, 380 TIL TCR clonotypes from sample BT21 were synthesised in pooled format, assembled, pooled and Nanopore sequenced to show the distribution of reads mapping to each TCR showing a tight distribution of TCRs with some dropouts. **d**, Percentage of error-free TCRs, as assessed number of Nanopore reads having fewer alignment errors than predicted by the Q score of the respective read. **e**, combined plot of c and d showing that TCR dropouts are the main cause of error. **f**, alignment errors in reads mapped almost entirely to the PCR amplified CDR3 regions (lowest 20 % quantile of TCRs filtered to remove cloned TCRs with insufficient reads for accurate assessment). **g**, Heatmap of 380 TCRs demonstrating correct assembly per fragment on an individual TCR basis.

As proof of concept, we manufactured 380 TCRs from our ‘BT21’ melanoma TIL sample. We adapted the demultiplexing protocol published by Subramanian et al. for use with the high-yield, low-GC bias KAPA HiFi polymerase to minimize primer off-target annealing and ensure selective, high-yield amplification (see Methods). We cleaned and concentration normalised the resulting PCR products using a liquid handling platform, and assembled each TCR with its appropriate V, C and backbone modules using a non-contact dispensing system. We pooled the Golden Gate assembled TCRs, transformed them into *E.coli*, and performed quality control on the resulting plasmid prep. Nanopore long read sequencing revealed a median of 80 % of reads resolving to a correctly assembled TCR (Figure 3a), with errors representing both dropouts (missing TCRs) and incorrectly paired CDR3 regions (Figure 3f,g) as a result of imperfect oligonucleotide demultiplexing. Overall, we found that high-throughput arrayed assembly of TCRs using makeTCR resulted in a tight distribution of TCR frequencies suitable for downstream functional screening.

### Simple software platform to facilitate TCR manufacture

To simplify and speed the process of cloning and validating TCRs we developed a graphical, web-based tool to automate the process of generating the oligonucleotide sequences required for modular TCR assembly (Figure 4a). The makeTCR software platform accepts diverse input formats, including the Adaptive Immune Receptor Repertoire standard (used by tools such as MiXCR ^25^), the *\*.consensus_annotations.csv* format generated by 10X Genomics’ Cell Ranger platform, and the output from VDJdb ^26^ (Figure 4b). Users can determine which TCRs should be cloned based on TCR frequency, or by submitting a list of target CDR3 sequences, with the software internally resolving ambiguous inputs to the IMGT format (e.g. “ASQY” to ”CASQYF”). The platform facilitates the cloning of both positive controls (HLA-matched TCRs from VDJdb or MR1-restricted TCRs that have been reported to show broad reactivity against tumor cell lines ^27^), and negative controls (randomly selected TCRs from the repertoire). Using the advanced options users can select to clone only TCR within subsets of interest, selecting between αβ, γδ, MAIT or CAIT TCRs (Supplemental Figure 5). Users can then select between the available makeTCR constant regions and expression vectors using annotated dropdown lists, and can easily integrate custom modules in a code-free manner by simply uploading a module’s sequence in fasta format. As a fraction of TCRs contain long homopolymer runs in the CDR3 regions that are both difficult to synthesise and accurately sequence (Figure 4d), makeTCR removes these by silent mutagenesis.

**Figure 4:**
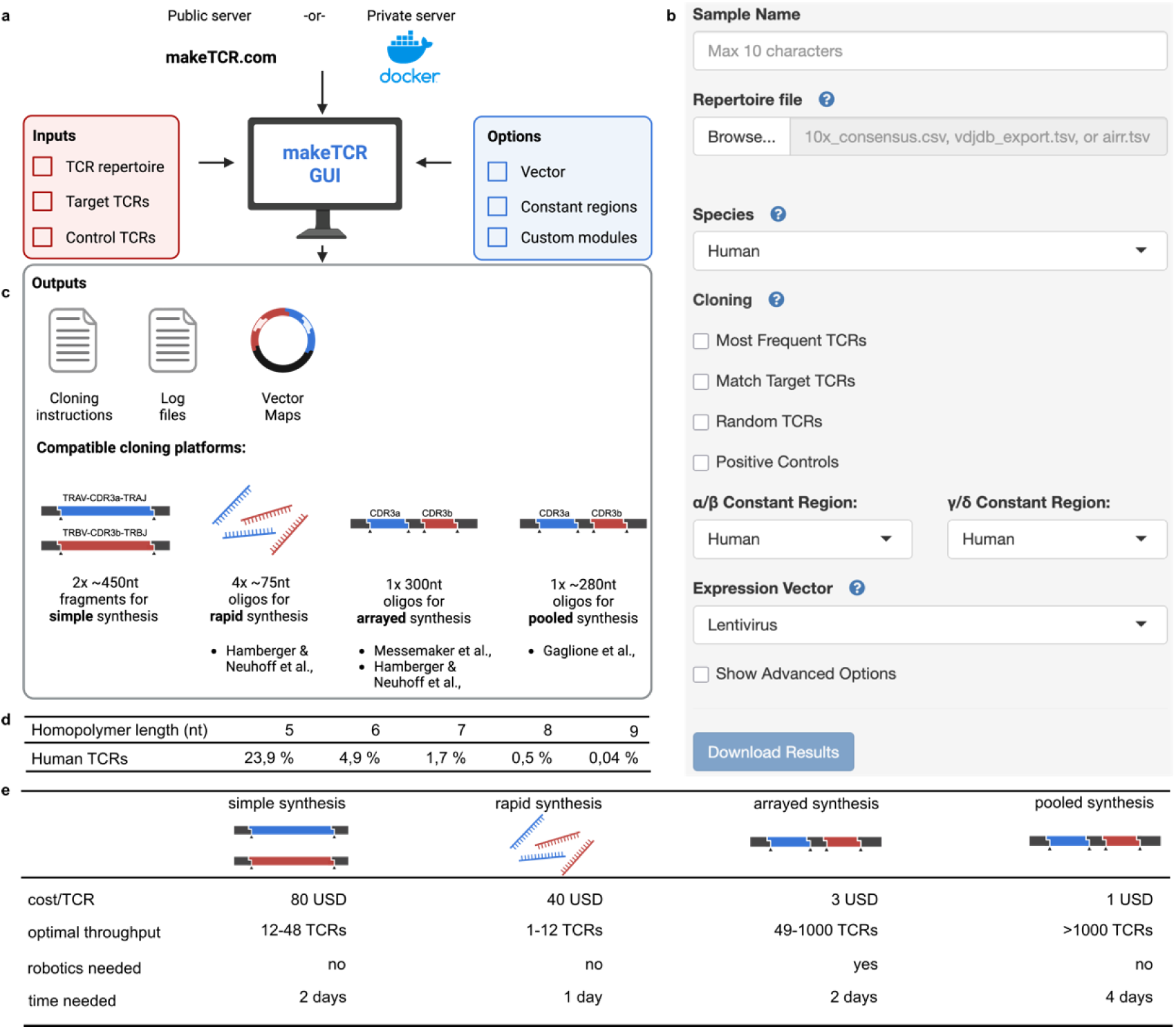
makeTCR graphical interface. **a**, schematic of the makeTCR software platform inputs and outputs. **b**, graphical interface of the makeTCR graphical software platform. **c**, Schematic of outputs provided by makeTCR. Besides log information and vector maps, cloning instructions for a range of cloning workflows are provided. **d**, CDR3-TR*J regions from TCRs derived from a human TIL repertoire natively contain nucleotide homopolymer runs which can be difficult to sequence; makeTCR depletes such homopolymers by silent mutagenesis. Listed are frequency of homopolymers in a human TCR repertoire. **e**, overview and comparison of cloning workflows supported by makeTCR; additional detail of oligo structures shown in Supplemental Figure 6).

Once the input parameters are set, the makeTCR software generates vector maps for each TCR (required for QC assemblies), cloning instructions (in both human readable and automation compatible formats), and the DNA sequences that must be ordered for each TCR (Figure 4c). A separate output folder is made for data relating to each method of TCR manufacture; simple synthesis (in which the entire VDJ sequence is exported for users who wish to clone TCRs without access to the pre-cloned modules from makeTCR or TCRAFT), rapid synthesis by oligo annealing, synthesis from arrayed demultiplexed oligonucleotide pools, and the TCRAFT pooled TCR assembly system ^21^(Figure 4e, Supplemental Figure 6). For arrayed TCR synthesis of more than 383 TCRs we implemented a variant of the two-step nested oligonucleotide demultiplexing strategy of Messemaker and colleagues ^28^ whereby TCRs are first demultiplexed into pools of 383 using an outer primer pair, and then further demultiplexed in individual wells using a standardised inner pair of primers. This approach allows for a standardised set of 384 primer pairs to be reused for each pool, simplifying setup by allowing replica primer plates to be stamped out. By adding filler DNA sequences to facilitate PCR cleanup, we created an oligonucleotide format that combines simple demultiplexing with flexible TCR assembly.

Lastly we extensively documented software options to assist users in selecting the most appropriate format for their TCRs, providing best practice guidelines and linking to the source scientific literature. The software platform is entirely open source, is hosted freely online at https://makeTCR.com, and can be run on private servers as a Docker container when TCR sequences cannot be publicly disclosed.

## Discussion

TCR based immunotherapies are of increasing interest due to the much broader range of targets available to TCRs relative to chimeric antigen receptors (CARs), the decreasing costs of transgenic cell manufacture ^29^, and the opportunities for developing truly personalised cell therapies ^3^. However, the TCR field has not yet coalesced around a rapid, high-fidelity synthesis solution that enables simple, low-cost, flexible yet scalable manufacture of TCRs. We addressed these limitations with the makeTCR platform, which we built to meet the needs of researchers working across different model systems and therapeutic contexts, enabling consistent and reliable TCR assembly of both αβ and γδ TCRs, for both human and mouse sequences. By utilising data-optimized Golden Gate assembly design techniques ^30^, makeTCR allows for versatile assemblies of VDJ regions grafted onto a TCR constant region of choice without compromising assembly fidelity. Furthermore, makeTCR enables standardized TCR generation regardless of downstream application by providing compatible vectors for all commonly used TCR expression systems; vectors that can be re-utilised for BsmBI mediated cloning of other molecular cargos. For the first time we demonstrate the utility of using Phi29-mediated RCA to substitute for traditional bacterial culture in propagating reusable cloning modules, representing a practical advance that reduces the labour and resources required to maintain a modular plasmid collection while retaining TCR assembly fidelity.

A key limitation of previous TCR synthesis techniques was the lack of a software platform to automate the process of generating TCRs from information contained in TCR repertoire datafiles. We developed a free, extensible, open-source graphical platform that allows researchers to upload repertoire sequencing data, select which TCRs they wish to clone, and receive all required DNA sequences, cloning instructions, and vector maps for performing quality control sequencing. By outputting TCR manufacturing solutions for all recently reported synthesis strategies, the makeTCR software platform represents a ‘one stop’ tool that covers the gamut of TCR manufacture from rapid flexible cloning to high-throughput screening. The software platform is available at https://makeTCR.com, can be run locally (in case of data privacy issues), and can easily be expanded in a code-free manner by users who wish to use novel modules – for example TR*C domains linked to suicide switches enabling the targeted depletion of TCR transgenic cells ^31^.

We have demonstrated that makeTCR enables rapid prototyping of TCRs: novel, patient-derived, tumor-reactive TCRs could be identified, manufactured and functionally validated within 48 hours – a process that previously took multiple weeks ^32^. This illustrates the potential of this approach for prioritising patient-derived TCRs with optimal properties such as dominant membrane expression for personalised clinical applications. Our results are particularly timely given the recent FDA ‘Fast Track’ designation for mRNA encoded TCRs as a monotherapy for HBV-related hepatocellular carcinoma (NCT05195294). This suggests that pipelines combining makeTCR’s rapid TCR assembly with cell-free, GMP grade, endotoxin-free manufacture of mRNA, DNA doggybones ^33^ or nS/MAR vectors ^34^ could further accelerate - and thereby reduce the cost of - personalised TCR therapies.

We have shown that makeTCR scales to high-throughput TCR manufacturing using arrayed TCR manufacture, and supports the design of oligos for pooled library cloning ^21^. These large numbers of TCRs can be leveraged in pooled functional screening applications, for example using synthetic reporters of T cell activation to uncover CDR3 motifs binding recurrent driver mutations ^35^. These screens pave the way for the generation of large curated datasets that can feed into motif discovery and next generation machine learning algorithms aimed at deciphering TCR antigen specificity and predicting therapeutic TCRs for specific diseases or patient subtypes. To facilitate such studies, and the use of the makeTCR resource by the broader immunology research community, all makeTCR modules and vectors are available from both Addgene and the European Plasmid Repository.

## Material availability

Human germline TRAV and TRBV plasmids, as well as a selection of TR*C plasmids and makeTCR compatible vectors, have been submitted as 96 well plate plasmid collection to Addgene (www.addgene.org) – see Supplemental Table 1, plate ‘2_Hs_V’.

Murine germline TRAV, TRBV, TRGV and TRDV plasmids, Human TRGV and TRDV, and an extensive collection of TR*C modules and expression vectors are available from the European Plasmid Repository (www.plasmids.eu) – see Supplemental Table 1, plates ‘3_Mm_V’ and ‘4_GD_Misc’.

## Code availability

The makeTCR service is available at https://makeTCR.com, and on publication the source code will be made freely available under MIT license at https://gitlab.com/EdGreen21/makeTCR.

## Acknowledgements

- We gratefully acknowledge the data storage service SDS@hd supported by the Ministry of Science, Research and the Arts Baden-Württemberg (MWK) and the German Research Foundation (DFG) through grant INST 35/1503-1 FUGG and INST 35/1503-1 FUGG as well as the German Cancer Research Center Next Generation Sequencing and Flow Cytometry Core Facilities.
- The German Ministry of Education and Science (National Center for Tumor Diseases Heidelberg NCT 3.0 program ‘Precision immunotherapy of brain tumors’ and the DKTK program) to MP (DKTK equipment grant).
- Federal Ministry of Education and Research (BMBF) and the Ministry of Science Baden-Württemberg, Excellence Strategy of the Federal and State Governments of Germany EMS, flagship ‘Engineering Molecular Systems’ spotlight proposal ‘Synthetic Immunology’ and ‘Sequencing Oligo Libraries’ (ExU 6.1.9).
- Hector Cancer Institute Seed Funding Program to MP, project 445549683,
- M-TN is supported by the Vector Stiftung (P2024-0750).
- CLT is supported by a DKFZ Hector Seed Funding Kick-start EarlyCareer fellowship (C-3PO).
- TB is supported by the Helmholtz-Institute for Translational Oncology Mainz (HI-TRON Mainz) as by a German Cancer Consortium DKTK, DKFZ Joint Funding project award (TRUST) to MP.
- MP received funding from the European Research Council (ERC) under the European Union’s Horizon Europe project 101141901.
- We gratefully acknowledge Dr Kim Boonekamp for help preparing figures.
- Figure panels created using BioRender.com.

## STAR Methods

### Design and testing of optimal Golden Gate overlaps for TCR assembly

Mutually orthogonal Golden Gate overlaps for assembling TCRs as polycistronic transcripts were iteratively designed to bracket the methionine at the start of TR*V genes, the Cysteine residue defining the CDR3 region, the first complete amino acid of TR*C, and either the end of a T2A linker (for TRBC-T2A and TRGC-T2A) or a TAA encoded stop codon (for TRAC and TRDC) with help of NEB’s web based Ligation Fidelity calculator ^30^ and targeted silent mutagenesis. The final overlaps are shown in Supplemental Figure 1.

### Golden Gate assembly fidelity assays using amilCP

We codon optimised the coral *Acropora millepora* chromoprotein amilCP for expression in *E.coli*, splitting the coding sequence into six modules linked by the same Golden Gate overlaps and flanked by BsmBI recognition sites as used to make an αβ TCR (CCAT, TGTG, GACC, GATG, CTGT, ATCC, TAAC). amilCP expression was driven by the constitutive bacterial promoter BBa_J23110 and terminated by the *E. coli* histidine operon terminator. The resulting sequence was ordered as a dsDNA fragment from Twist Biosciences and cloned into a custom backbone. This modular amilCP construct is used as a positive control for makeTCR assemblies; full construct sequence details are available from Addgene (plasmid #233942). The amilCP modular control vector was prepared as miniprepped plasmid, and this was further amplified by RCA and assembled into the makeTCR third generation lentiviral backbone in a BsmBI mediated Golden Gate reaction according to the manufacture’s guidelines (plasmid #233943). Assemblies were transformed into NEB stable bacteria, plated on agar and grown overnight at either 30°C or 37°C before allowing the chromoprotein to mature at room temperature (n=4 independent assemblies per condition, 3 technical replicates per assembly).

### Manufacture of BsmBI site flanked germline V gene modules

Germline V gene sequences were domesticated by using silent mutagenesis to remove extraneous binding sites for the IIs restriction enzyme BsmBI (CGTCTC) required for TCR assembly using custom R scripts. To ensure maximal downstream compatibility V genes were also optimised to deplete the IIs enzyme BsaI (GGTCTC), the nickase Nt.BbvCI sequence (CCTCAGC), and the rare cutter SmiI/SwaI (ATTTAAAT) to allow for post Golden Gate assembly linearisation of unassembled input material. Sequences were ordered as dsDNA fragments from either Twist Biosciences or Integrated DNA Technologies (IDT) and assembled into the ‘pMC’ custom backbone vector by BsaI mediated Golden Gate assembly. Full vector sequences are available in Supplemental Table 1, as well as from Addgene and the European Plasmid Repository.

### RCA mediated amplification of V gene modules

1 ng of each germline V gene plasmid was amplified according to the manufacturer’s instructions by randomly primed RCA using New England Biolab’s phi29-XT RCA kit in a total volume of 20 µl. Prior to bead purification reactions were diluted 1:5 with nuclease free water to avoid clumping of high molecular weight DNA in the subsequent magnetic bead cleanup (AMPureXP Beckman Coulter). Beads were added to the reaction (0.6X bead to reaction volume) and purification was completed according to the manufacturer’s instructions.

### Rapid TCR cloning using annealed oligos

Oligos encoding the CDR3-TR*J region were synthesised with a 5’ phosphate moiety using a SYNTAX device (DNA Script SAS). Forward and reverse oligos encoding a CDR3-TR*J fragment were annealed in nuclease-free Duplex Buffer (30 mM HEPES, pH 7.5; 100 mM potassium acetate, IDT) in a thermocycler (95°C for 2 min then cooled to 25°C over 25 mins) to generate 2 µM annealed dsDNA with cohesive ends compatible with the respective 4 nt makeTCR TRAV/TRAC or TRBV/TRBC assembly sites. Annealed oligos were diluted to 0.2 µM in nuclease-free water and then used in Golden Gate assembly reactions at twice the molar ratio of other components (0.1 pmol of each annealed oligo pair, 0.05 pmol TR*V and TR*C, 0.034 pmol precut vector backbone). Due to the risk of melting annealed oligos, the Golden Gate assembly protocol was modified to include an initial low temperate step (25°C for 20 min), followed by 30 cycles of 42°C/16°C for 1 min each, then an additional 15 min at 42°C, and a final heat inactivation step at 60°C for 5 min.

### Generation in vitro-transcribed TCR mRNA constructs

For in vitro transcription, TCRs were assembled into the makeTCR lentiviral backbone which includes an optimised T7 promoter downstream of the elongation factor one alpha (EF1α) promoter used to drive expression in transduced T cells ^37^. A linear template for IVT reactions was generated by PCR utilising 0.2 µl of the Golden Gate TCR assembly reaction in a 20 µl reaction using SuperfiII polymerase (Thermo Scientific). 20 cycles of PCR were performed using a Tm of 66°C using the forward primer CAGGTGTCGTGACGTTTAGTG and the reverse primer GGCTGGCACGAAATTGTTG. The resulting PCR product was cleaned using AmpureXP beads (0.6X ratio) and used as a template for the T7 mScript Standard mRNA Production System (CELLSCRIPT C-MSC11610). mRNA was m7G capped and enzymatically polyadenylated following the manufacturer’s instructions.

### Quality control of TCRs generated from annealed oligos

IVT templates for individually assembled TCRs were prepared as barcoded libraries compatible with Nanopore sequencing using the Native Barcoding kit (SQK-NBD114.24, Oxford Nanopore Technologies). Libraries were sequenced on Flongle flow cell (R10.4.1) on a MinION Mk1D (Oxford Nanopore Technologies) for 24 h. Reads were demultiplexed and basecalled using the Dorado basecaller implemented within MinKNOW, and then aligned to the appropriate TCR reference sequencing using the Clara Parabricks (v4.3.2-1) implementation of Minimap2 (v2.26) ^38^ . The Dorado basecaller assigns a quality (q-score) to each read; if a read aligned to the reference sequence with fewer mismatches than would be expected from the q-score then that read was called to be error-free.

### Isolation and expansion of REP T cells from healthy donor PBMCs

PBMCs from healthy donors were isolated from heparinized blood. 15.5 ml of Ficoll Paque Plus Media (Cytiva) was loaded per Leucosep tube (Greiner Bio-One). After adding 3 ml of PBS (Sigma), up to 25 ml of blood was loaded on top and a density-gradient centrifugation was performed at 800 g. After collection of the interphase, PBMCs were washed twice with PBS and frozen in a controlled rate freezing device at −80 °C in 50 % freezing medium A (60 % X-Vivo 20 and 40 % fetal calf serum) and 50 % medium B (80 % fetal calf serum and 20 % dimethylsulfoxide). Cells were stored in liquid nitrogen at −140 °C until further analysis.

The rapid expansion protocol was used to expand T cells. PBMCs from three independent donors were irradiated at 40 Gy using a Gammacell 1000 (AECL) radiation device to serve as feeder cells. Then, 1 × 10^7^ cells from each donor were pooled together, cells were spun down (400 g, 10 min, room temperature) and resuspended in rapid expansion media (X-Vivo15 (Lonza, BE02-060Q), 2% human AB serum (H4522-100ML, Sigma-Aldrich), 2.5 µg/ml Fungizone (15290-018, Gibco), 20 µg/ml gentamicin (2475.1, Roth), 100 IU/ml penicillin and 100 µg/ml streptomycin (15140122, Life Technologies)). Next, 150,000 PBMCs were plated into a standing T25 flask and 666 ng of OKT-3 antibody (Life Technologies, 16-0037-85) was added to the culture and the flask was topped up to a total volume of 20 ml. The next day, 5 ml of X-Vivo15 supplemented with 2 % AB serum containing 7,500 IU interleukin-2 (IL-2) was added to the culture. Three days later, 12.5 ml of medium was removed and replaced with 12.5 ml of X-Vivo15 supplemented with 2% AB serum containing 600 IU ml−1 IL-2. We estimated the REP cultures to contain between 40 and 50 % CD8+ T cells using flow cytometry.

### TCR reactivity screening via flow cytometry

TCR-encoding RNA was electroporated into expanded healthy donor PBMCs using the Lonza 4D-Nucleofector (program EO-115, solution P3 supplemented according to the manufacturer’s recommendations). Transfected cells were plated in 48-well plates containing TexMACS media (130-097-196, Miltenyi) supplemented with 2% human AB serum. At 18–24 h after electroporation, cells were collected and 50 IU ml^−1^ benzonase (YCP1200-50KU, Speed BioSystems) was added to avoid cell clumping. TCR expression levels were measured via flow cytometry with markers including fixable viability dye (eFluor 450, eBioscience), CD3 (clone HIT3A, BUV510, BD), and mTCRb (clone H57-597, PE, Biolegend).

To assess TCR reactivity, a total of 150,000 T cells and 75,000 cells of the patient-autologous tumor cell line were cocultured in U-bottom 96-well plates in a total volume of 200 µL for 4 hours. Wells with only T cells, or T cells and TransAct beads (130-111-160, Miltenyi) were used as negative and positive controls, respectively. Markers included fixable viability dye (eFluor 450, 1:1,000 dilution, eBioscience), CD3 (clone UCHT-1, BUV395, 1:100 dilution, BD), CD69 (Clone FN50, PE-Cy7, 1:200, Biolegend) and mTCRb (clone H57-597, 1:50 dilution, PE, Biolegend). Samples were acquired on a Biorad ZE5 and flow cytometry data were analyzed using FlowJo software, v10.6.2 (FlowJo LLC).

TCRs were classified as reactive or nonreactive based on flow cytometry data acquired after coculture. The percentage of CD69 (%CD69) was quantified by gating on viable CD3^+^ singlets. TCRs were included in the analysis if the mTCRβ expression was >10%. The CD69 signal per TCR after coculture with the cell line (‘TCR versus cell line’) or after running the coculture assay without stimulation (‘TCR, unstimulated’) was corrected for background by calculating (%CD69 TCR vs cellline − %CD69 TCR, unstimulated) − (%CD69 Mock vs cellline − %CD69 Mock, unstimulated) where mock refers to expanded T cells electroporated without TCR-encoding RNA. TCRs were classified as reactive if the background corrected %CD69 signal per TCR was larger than 2× the standard deviation of the %CD69 signal measured in all samples without stimulation. Where a TCR clonotype expressed two α chains, data are presented for the α chain resulting in the higher %CD69 expression (i.e. the functional pair).

### ssDNA oligo pool design

Sequences encoding the CDR3a-TRAJ and CDR3b-TRBJ regions for each TCR were modified by silent mutagenesis to remove BsmBI, PaqCI and SwaI restriction enzyme sites. In cases were TCR sequences were to be analysed by nanopore long read sequencing data, homopolymer runs of 6 nucleotides or more were removed by further silent mutagenesis to prevent sequencing artifacts around these regions. BsmBI sites were then appended to the end of each CDR3-TR*J sequence, and filler DNA added to pad all sequences to 210 nucleotides using custom R code to add GC balanced random bases without introducing additional restriction sites, homopolymers, or primer binding sites. Unique pairs of 20nt long primers from Subramanian et al ^24^ were then added to the ends of the design to make a total of 250 nucleotides. An example sequence is shown in Supplemental Figure 4; this strategy allowed for cloning up to a combined total of 61 amino acids for CDR3a-TRAJ and CDR3b-TRBJ.

### Library demultiplexing PCR setup

Library demultiplexing was performed in 15 µL reactions using KAPA HiFi HotStart ReadyMix (Roche) with 0.05 ng of single-stranded DNA (ssDNA) oligo pool per well in a 384-well plate format. The demultiplexing PCR mastermix was dispensed into 384-well plates using a microfluidics based non-contact dispenser (Formulatrix Mantis). Two orthogonal primers were dispensed into each well at a final concentration of 0.2 µM, directly from 100 µM 96-well stock plates, using a 96-well plate based non-contact liquid handler (Dispendix I.DOT). We found that cooled PCR setup was essential to prevent KAPA HiFi polymerase’s 3’-5’ exonuclease activity truncating primers. Following reaction setup, plates were centrifuged at 1,000 rcf for 1 min and subjected to orbital shaking on a Bioshake iQ at 3,000 rpm for 3 min. Plates were centrifuged again prior to PCR amplification in an Eppendorf x50t under the following modified conditions: an initial three-step touchdown protocol starting at 65°C was implemented to enhance amplification specificity, followed by 33 cycles of PCR with an annealing temperature of 62°C and an extended annealing time of 40 sec to promote specific priming and efficient on-target amplification despite elevated annealing temperature. We found that using this high primer annealing temperature (relative to the original publication) was key to ensuring specific amplification of target oligos.

Post-PCR cleanup was performed using AMPure XP beads, dispensed at a 1:1 bead-to-reaction volume ratio using the Formulatrix Mantis. Plates were then transferred to an Agilent Bravo liquid handling system to perform a magnetic bead-based DNA purification in 384-well format. PCR products were eluted in Greiner UV Star plates and post-cleanup DNA concentrations were measured via absorbance (Varioska LUX Multimode Microplate Reader); custom scripts were then used to normalise all wells to a uniform concentration of 1 ng/µL in deep-well plates.

### Golden Gate mediated arrayed assembly of thousands of TCRs

Golden Gate assembly of T cell receptors (TCRs) was performed using the NEBridge® Golden Gate Assembly Kit (BsmBI-v2) following the manufacturer’s recommendations, scaled down to a reaction volume of 2 µL. Assembly reactions were prepared in 384-well plates using a Dispendix I.DOT non-contact dispenser to give a ratio of 1 CDR3 oligo : 0.91 TR*V : 0.9 TR*C : 0.67 Vector. Corresponding αβ V gene segments, preset constant region modules, and the respective vector of choice were automatically dispensed onto 1 ng of purified oligo DNA per well. Following reagent dispensing, plates were mixed on a Bioshake iQ at 3,000 rpm for 3 min and centrifuged at 1,000 rcf for 1 min to collect the reaction. Assembly was performed in an Eppendorf X50t thermal cycler using a custom temperature profile: reactions were incubated at 42°C for 30 min, followed by 30 cycles of 42°C/16°C for 1 min each, then an additional 30 min at 42°C, and a final heat inactivation step at 60°C for 5 min.

### TCR library preparation

To generate the pooled plasmid library, 2 µl of each assembled TCR was combined into a single well using an automated liquid handling system (Bravo, Agilent Technologies). Prior to bacterial transformation, the pooled assemblies were purified using AMPure XP beads (Beckman Coulter) at a bead-to-sample ratio of 0.8X to enhance electroporation efficiency. Assembled plasmids were eluted in 60 µl of nuclease-free water and subsequently transformed into electrocompetent *E. coli* (NEB 10-beta, New England Biolabs) following the manufacturer’s protocol. To minimize recombination within lentiviral long terminal repeats, bacterial outgrowth was performed at 30 °C for 1 h in NEB’s proprietary recovery medium. Cultures were then transferred directly into selective LB medium supplemented with 100 µg/mL carbenicillin. To assess transformation efficiency and estimate library coverage, 10 µL of the outgrowth was plated on 10 cm LB-agar plates containing the same antibiotic. For all transformations, a minimum of 500X library coverage was maintained. Cultures were expanded at 30 °C for 16–24 h prior to plasmid extraction using a ZymoPURE II Plasmid Maxiprep Kit (Zymo Research).

### Nanopore sequencing of TCR libraries

0.01 ng of generated plasmid libraries were used as input for amplicon generation to add UMI adapters to the end of constructs as described by Amstler et al., 2024. UMIs were incorporated by two cyles of PCR (2 min extension at 72°C, 30 s annealing at 60°C) with Platinum SuperFi II PCR Master Mix (Invitrogen) using 0.01 ng of the prepared TCR library and 0.5 nM of primers GTCTCTGCTCGACTAACCACTTTVVVVVTVTTVVVVTTVVVTTGTGTGGCTGGCACGAAATTG and GCTCTCATACGAACTCGTCCTTTVVVVVTTVVVTTVVVVTTVVVVTTTAAGTGCAGTAGTCGCCGTG (IDT, machine mixed random bases incl. PAGE-purification. This amplicon was then purified with magnetic beads (Ampure-XP Beckman Coulter, 0.8x ratio). Subsequently, the UMI-tagged product was subjected to a second PCR using Platinum SuperfiII polymerase (Invitrogen) with universal primers binding the last 20 bases of both 5’ adaptors of the UMI primers. 0.5 nM of either primer (GCTCTCATACGAACTCGTCC, GTCTCTGCTCGACTAACCAC) was used to amplify the product derived from PCR1 for 25 cycles (annealing 60 °C, 10 s, extension 72 °C 60 s). Nanopore compatible libraries were then generated using either the Ligation sequencing kit (SQK-LSK114) or the Native Barcoding kit (SQK-NBD114.24, Oxford Nanopore Technologies) and sequenced on a PromethION 2 Solo device for three days using a PromethION Flow Cell R10.4.1 or a flongle (R10.4.1) on a Mk1D for 24 h. To reduce compute time we implemented Clara Parabricks (v4.3.2-1) to run Minimap alignments on CUDA GPUs^38^. Secondary and supplementary alignments were subsequently filtered from the generated bam file using samtools ^40^.

### Calculation of correct assembly rate

The assembly rate was calculated as follows: when sequencing pooled, high-throughput manufactured TCRs amplicon reads were first anchored to a TCR reference by filtering for reads containing a perfect alignment to the first and last ten nucleotides of a TCR (thereby depleting reads mapping to residual empty vector that could not be assigned to any one TCR assembly). When sequencing barcoded TCRs generated by rapid TCR assembly this filtering was not required as reads could be mapped to the correct reference TCR by the barcode (derived from the SQK-NBD114.24 kit). Phred scores of the aligned fraction of reads were converted to error probabilities (10^(-Q_score/10); as described by Edgar and Flyvbjerg ^41^ the expected error rate for each read was characterized by the sum of error probabilities of the alignment region to the reference rounded to the nearest lower integer. The expected error rate was compared to the NM tag of the alignment allowing an estimation of the edit distance between a respective read and its cognate reference (observed errors within the read). Reads with fewer observed errors than expected on the basis of read quality were classified as a perfect assembly; the ratio of perfect assemblies to the number of reads aligning to a particular reference represents the assembly rate.

### Assessment of cloning fidelity

When sequencing pooled, high-throughput manufactured TCRs amplicon reads were first aligned against all full-length TCR reference sequences to establish the ground truth TCR for each read. When sequencing barcoded TCRs generated by rapid TCR assembly this filtering was not required as reads could be mapped to the correct reference TCR by the barcode (derived from the SQK-NBD114.24 kit). In both cases reads were then aligned to individual TCR sub-sequences (TRAV, TRBV, CDR3a, CDR3b, TRAC, and TRBC-T2A). Read IDs and their respective reference assignments were extracted from the BAM file, merged, and then compared to evaluate cloning fidelity for each reference. For each individual segment, the percentage of on-target reads aligning to the correct reference was determined (i.e. if a TCR utilised TRBV10-1, what % of reads for this TCR mapped to TRBV10-1).

## Supplementary Information

**Supplemental Figure 1:**
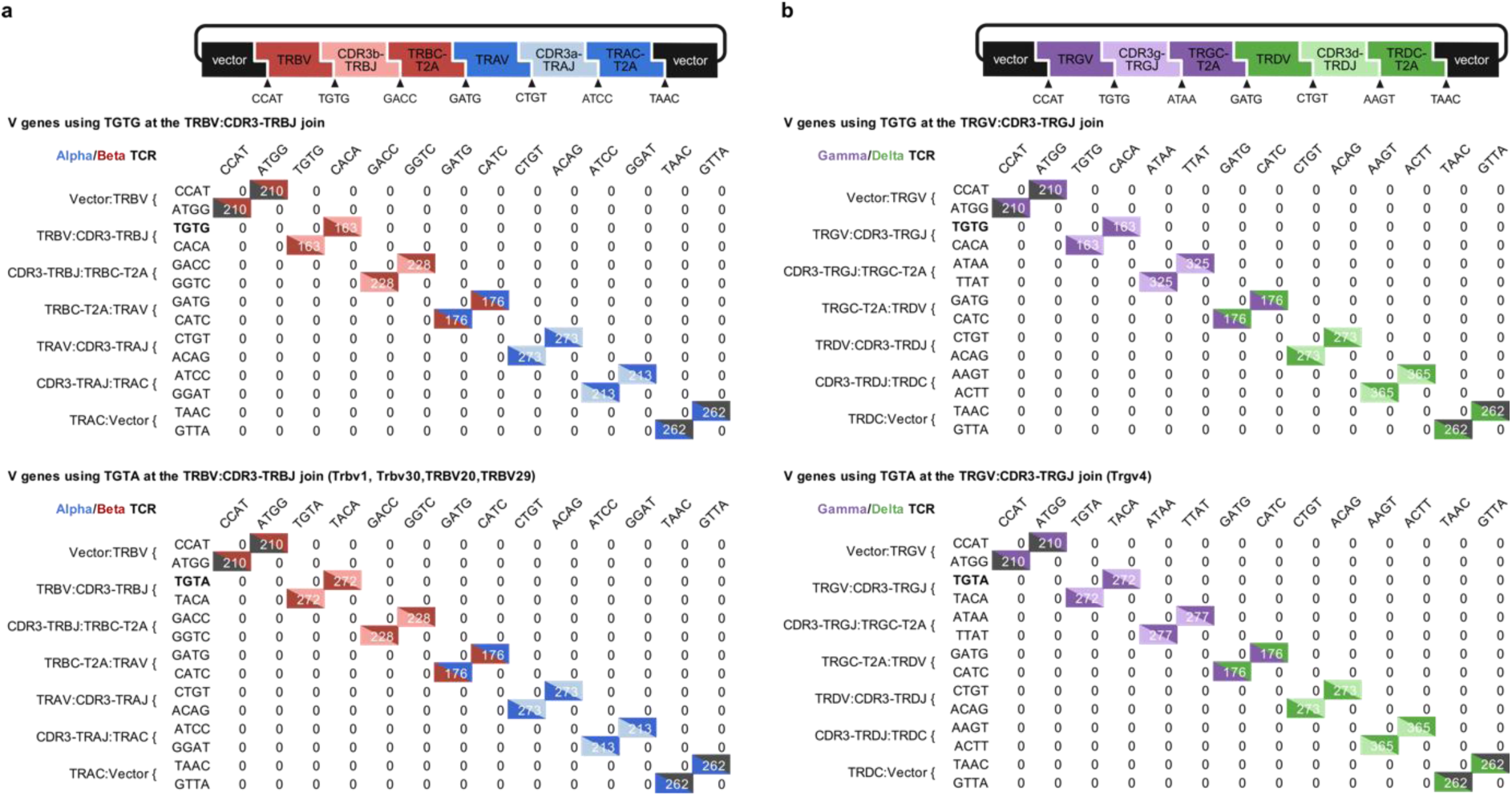
Predicted makeTCR Golden Gate Module Assembly Fidelities. **a;b**, Golden Gate fidelity matrices for cohesive ends linking makeTCR modules for both Alpha/Beta and Gamma/Delta TCRs.

**Supplemental Figure 2:**
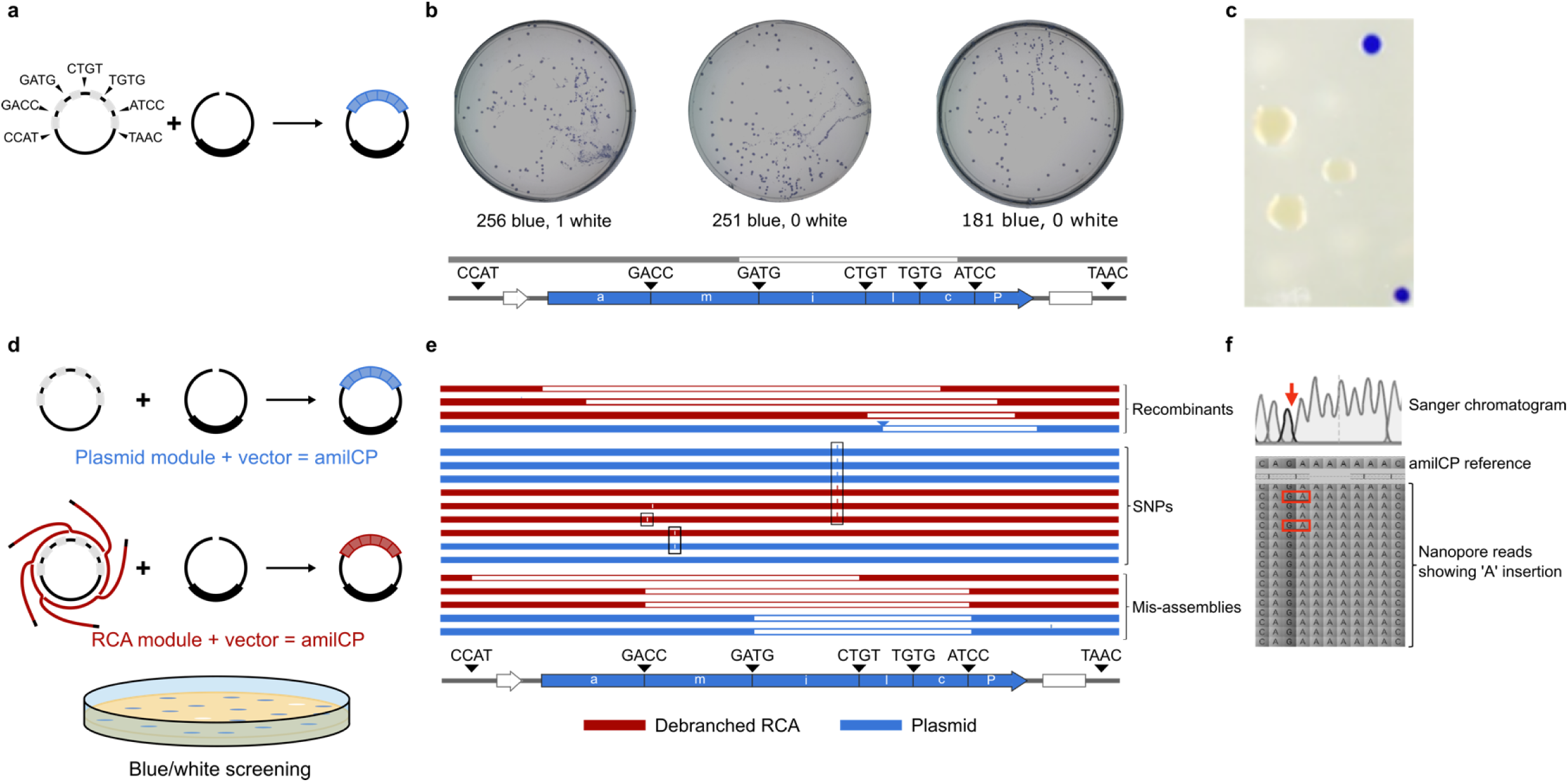
Validating assembly fidelity. **a**, A bacterial codon optimised expression cassette encoding the blue chromophore amilCP was split into BsmBI flanked modules linked by the cohesive ends selected to assemble an alpha/beta TCR (CCAT, TGTG, GACC, GATG, CTGT, ATCC, TAAC). **b**, The amilCP chromoprotein was assembled in a Golden Gate reaction and used to transform *E. coli*, resulting in 688 blue colonies and 1 white colony: an assembly fidelity of 99.9% vs the computationally predicted fidelity of 100%). The single white colony was Sanger sequenced to show a misassembly event joining the cohesive ends GATG and TGTG, resulting in a truncated non-functional chromophore. **c**, We confirmed previous reports that expression of amilCP from high-copy plasmids imposes at least a 25 % fitness cost on *E. coli* hosts (small blue colonies vs large white colonies)^15^; this ensured that our functional estimates of Golden Gate assembly fidelity were unlikely to be over-estimated due to the selective advantage of loss of amilCP expression in bacteria. **d;e**, Re-assembly of amilCP modules from a plasmid encoded (blue) or RCA amplified (red) template resulted in equivalent numbers of white colonies (Figure 1f-h). We sequenced white colonies assembled from either template types we found equivalent types of errors: amilCP truncation by bacterial recombination (sequence loss not matching module junction sites), Golden Gate mis-assemblies (deletion of regions exactly matching module junction sites), and single nucleotide polymorphisms (SNPs, raised lines representing nucleotide insertions, gaps representing missing nucleotide). We found that in almost all cases SNPs resulted in premature stop codons (black boxes), confirming the selective pressure against amilCP expression (**c**). **f**, We reasoned that SNPs shared between the RCA and plasmid templates must be present in the input plasmid material at levels undetectable by commercial Sanger and Nanopore plasmid QC sequencing. Manual examination of Nanopore plasmid sequencing reads showed that the most common SNP in both assemblies was indeed a rare variant present in the input material.

**Supplemental Figure 3:**
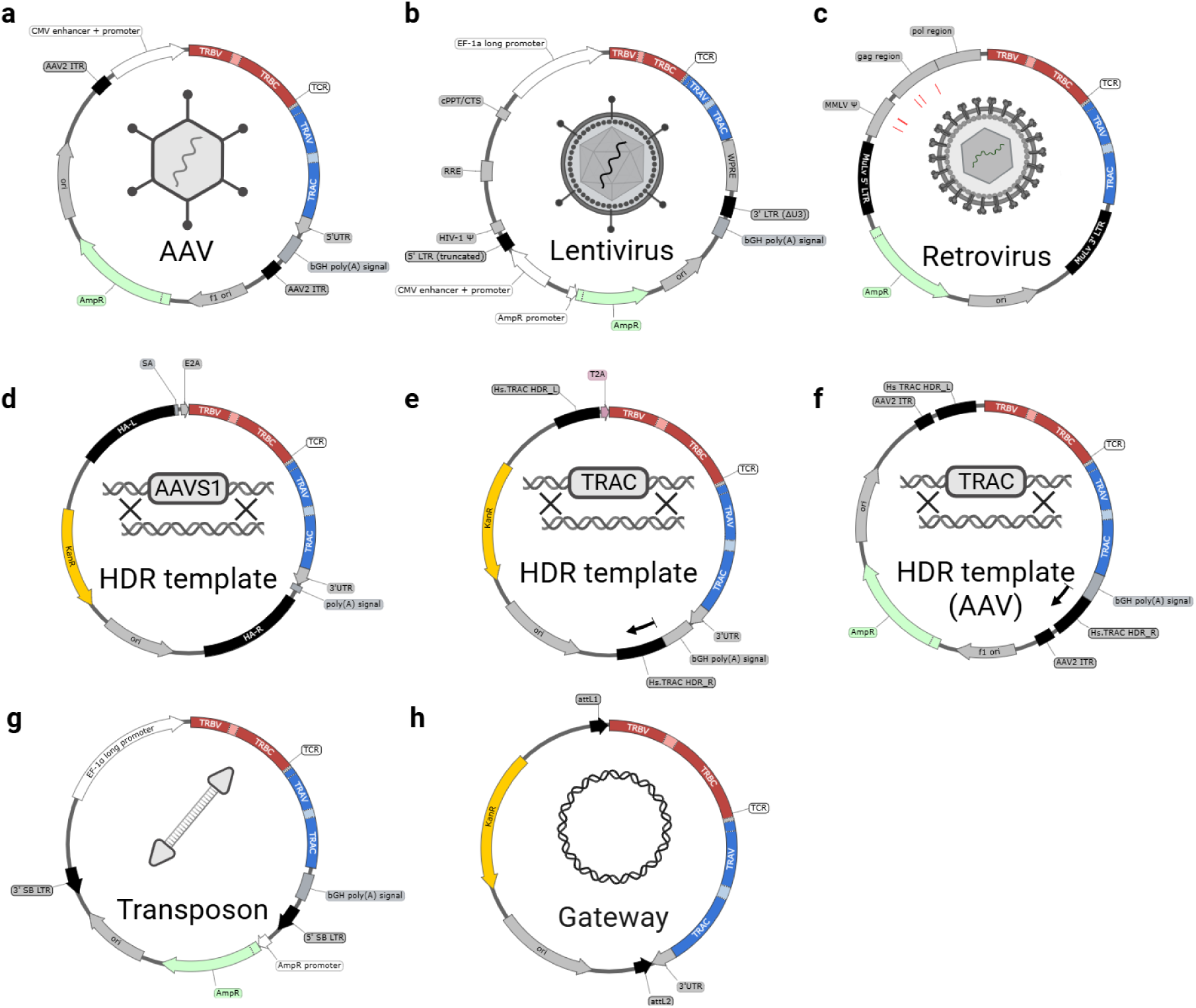
makeTCR expression vectors. **a**, AAV vector driving TCR expression using a CMV promoter. **b,** Lentiviral transfer vector driving TCR expression using a long EF1a promoter; truncated LTRs satisfy higher biosafety requirements. **c,** Retroviral TCR expression vector based on pMXS backbone using MMLV LTR region to express TCR. **d,** Vector for HDR mediated integration into the Human genome at the AAVS1; TCR driven by endogenous promoter. **e,** Vector for HDR mediated integration into the Human genome at the TRAC locus; transgenic TCR driven by endogenous TRAC promoter. **f**, AAV2 packaging compatible variant of **e**. **g**, Sleeping Beauty transposon vector; TCR expression driven by long EF1a promoter. **h**, Gateway^TM^ pENTR compatible vector for shuttling TCRs to existing Gateway^TM^ backbones.

**Supplemental Figure 4:**
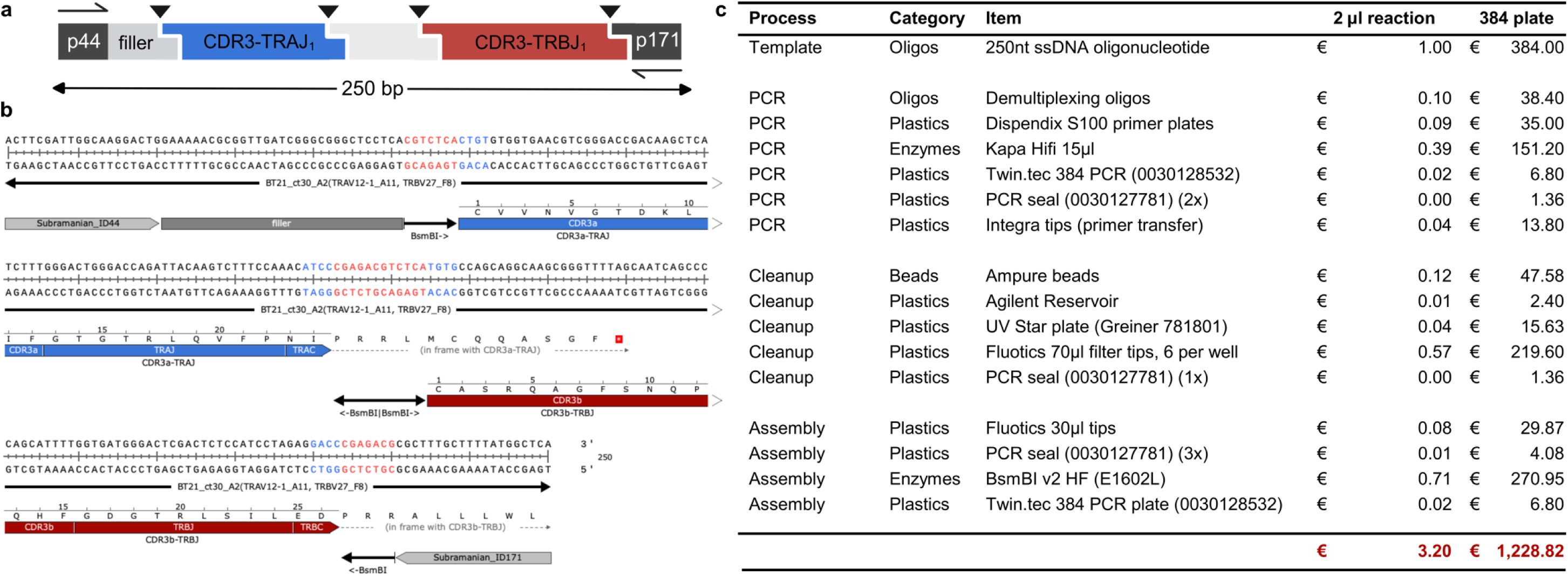
Example of a high-throughput makeTCR oligonucleotide and pricing breakdown. **a**, schematic of a 250nt oligo encoding makeTCR compatible CDR3-TRAJ and CDR3-TRBJ regions. **b,** nucleotide sequence of an example oligo sequence (shown as a dsDNA post PCR amplification product, DNA sequence encoding BsmBI sites highlighted in red, cohesive ends for ligation highlighted in blue). **c**, costs of high-throughput makeTCR assemblies when generating 2000 TCRs per manufacturing run.

**Supplemental Figure 5:**
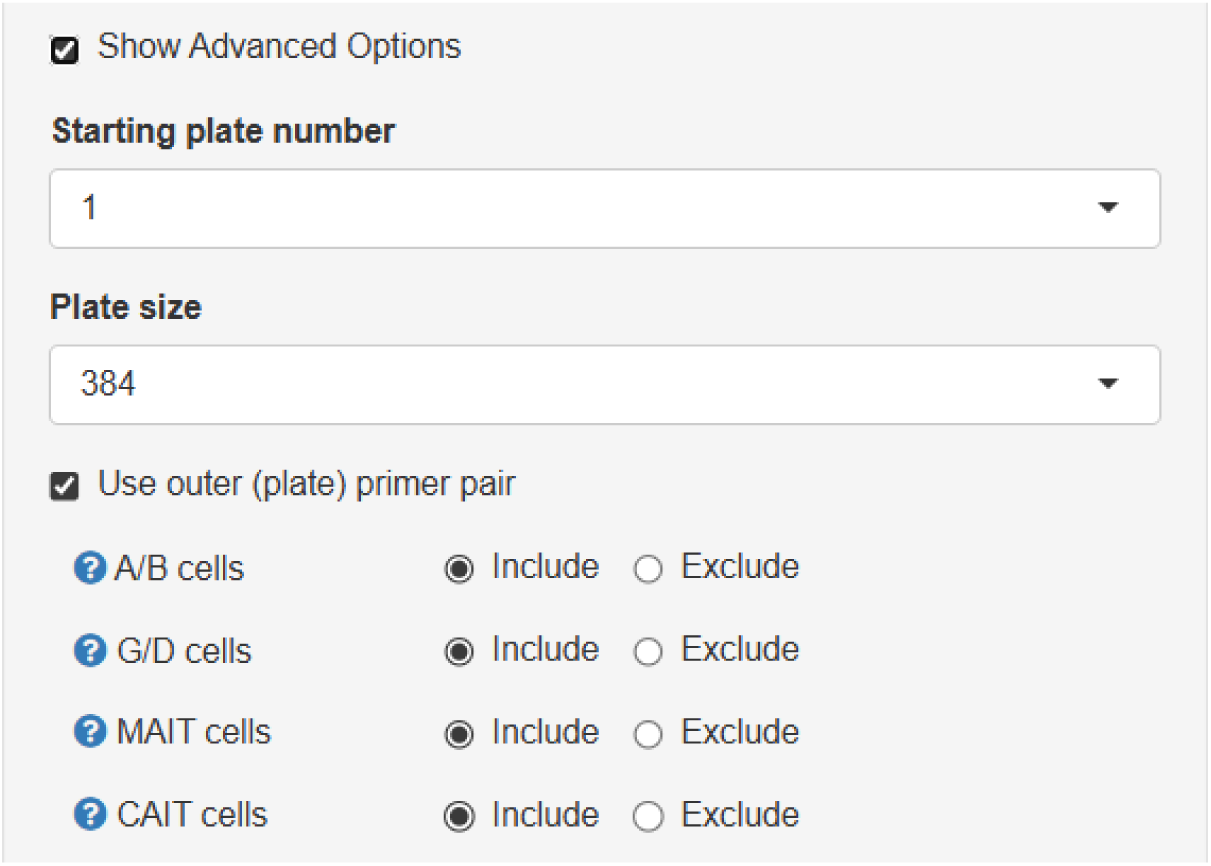
makeTCR platform advanced options. The makeTCR software platform allows users to select which pair of outer primers (plates 1-8) to append to TCRs, following the schema of Messemaker et al. ^28^. If more than 383 TCRs are selected for cloning, primers will be assigned in sequential plate order. Users can also configure whether they have access to 384 or 96 well liquid handlers, which modifies the automation script output and the naming schema for the vector reference maps to make it easy to track which well contains which TCR. Users can also select to only clone certain subtypes of TCR, selecting from αβ, γδ, MAIT and CAIT TCRs.

**Supplemental Figure 6:**
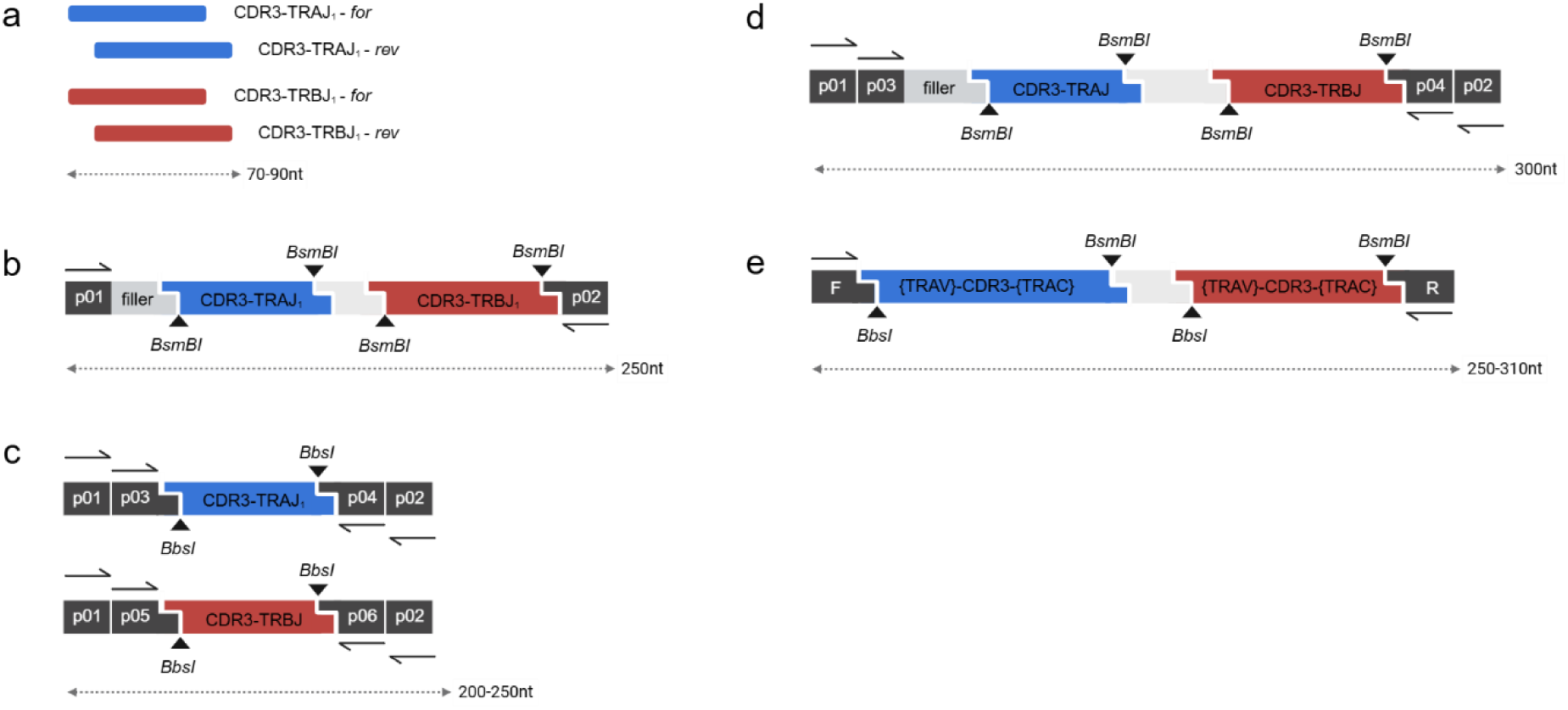
Schematic of TCR cloning oligo formats. **a**, makeTCR rapid format: four ssDNA oligos are annealed and used in modular assembly. Strategy requires 4x ∼80nt oligos per TCR. **b**, makeTCR arrayed oligo format: each oligo encodes both the alpha and beta CDR3 information, and is amplified from a pool using a unique pair of demultiplexing primers. A filler sequencer normalises the length of each oligo. Strategy requires 1x 250nt oligo per TCR. **c**, The original format proposed by Messemaker et al. in which pooled oligos are first amplified as pools using a plate-specific pair of outer enrichment primers (p01 + p02), before being demultiplexed using a well-specific primer pair (p03 + p04 or p05 + p06). Alpha and beta CDR3 information is on separate oligos. Strategy requires 2 oligos per TCR, ranging in length from 200-250nt. **d**, makeTCR implementation of the Messemaker et al. two round demultiplexing strategy, with the addition of a filler sequence to normalise the length of all oligos. Strategy requires 1x 300nt oligo per TCR. **e**, The TCRAFT pooled oligo assembly strategy in which 97% of TCR clonotypes can be encoded on a single oligo (1x 300nt/TCR), and the remaining 3% of long CDR3 TCR clonotypes are ordered as dsDNA fragments.

**Supplemental Table 1:** makeTCR Germline V Module Sequences. makeTCR modules are distributed on three 96 well plates: ‘p2_Hs_V’ containing the Human TRAV and TRBV germline modules, ‘p 3_Mm_V’ containing the Trav and Trbv modules for cloning Murine TCRs, and ‘p 4_GD_misc’ containing the modules for Human and Mouse gamma/delta TCRs, as well as additional TR*C modules, vectors and uncommonly used Murine germline Trav modules. Addgene_id = Addgene plasmid number, EPR_id = European Plasmid Repository plasmid number *(included as separate file due to table size)*.

**Supplemental Table 2:**
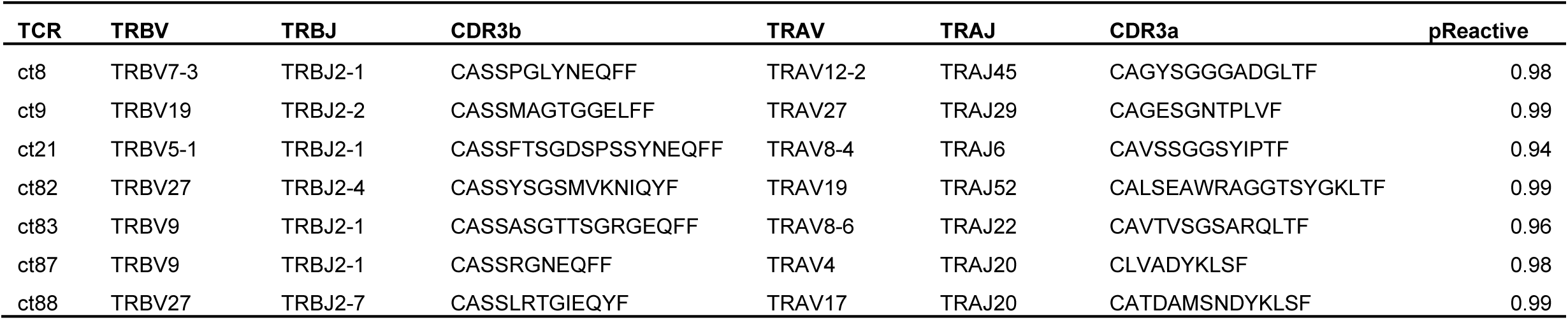
BT21 TCR sequences. Sequence of TIL derived TCRs tested in this study from the melanoma brain tumor metastasis sample ‘BT21’ previously described by Tan et al.^10^.. pReactive is the predicTCR calculated probability of a given TCR clonotype being tumor-reactive.

